# Binomial order is a speech marker of psychosis and thought disorder

**DOI:** 10.1101/2025.03.31.646413

**Authors:** Alice Luo, Jintian Luo, Michael Murphy

**Author notes:** Corresponding author: Michael Murphy. Author contributions: A.L. analyzed data and wrote the manuscript text. J.L. collected data. M.M. analyzed data, made figures, and wrote the manuscript text.

## Abstract

Thought disorder, characterized by disruptions in syntactic and semantic elements in language, is a core symptom of psychotic disorders. Understanding this language impairment is key to uncovering the underlying neuropathology and predicting treatment outcomes for individuals with schizophrenia and other psychotic disorders. Binomial ordering preferences (e.g. “salt and pepper” instead of “pepper and salt”), may be a quantifiable correlate of thought disorder and underlying linguistic impairments. We tested whether atypical binomial ordering can serve as a linguistic marker for psychosis symptoms. Participants with early-stage psychotic disorders and controls were recruited, and video-recorded interviews were transcribed for analysis. Identified binomial pairs were assessed using both the Google N-gram database and a logistic regression model to determine ordering preferences. Results showed that while both psychotic participants and controls preferred conventional binomial orderings, participants with psychotic disorders exhibited a higher rate of atypical binomial orderings. The use of atypical orderings was correlated with thought disorder, but not with other psychiatric symptoms or medications. Tracking binomial ordering can be a valuable marker of thought disorder but future studies are needed to determine whether this link remains stable or if it changes with disease progression.

## Introduction

Schizophrenia is a multifaceted psychiatric disorder that affects various cognitive and functional domains. Thought disorder is a core symptom of schizophrenia and can be described as disruptions in the organization, coherence, and logical flow of thought(1–3). While its presentation can vary across different individuals and different contexts, thought disorder can be broadly categorized as positive and negative(1,4,5). Positive thought disorder includes symptoms of tangentiality and derailment, while negative thought disorder indicates poverty of speech and poverty of content(1,6). Gooding et al. conducted a longitudinal study of thought disorder in the offspring of parents diagnosed with schizophrenia, affective disorders, or no psychiatric disorders(7). They found that the presence of subsyndromal levels of positive thought disorder was predictive of the development of affective and non-affective psychosis while the presence of subsyndromal levels of negative thought disorder was predictive of schizophrenia specifically, regardless of family history. This study highlights the potential of subsyndromal thought disorder as an early indicator of psychosis development and target early intervention. The presence and severity of thought disorder is associated with treatment outcomes, with more severe thought disorder predicting poorer response to interventions(8). Identifying and quantifying positive and negative thought disorder is crucial for informing therapeutic interventions for individuals at risk of developing psychotic disorders and for determining the prognosis for those already diagnosed.

Understanding language impairments is critical for diagnosing and elucidating the underlying neuropathology of schizophrenia and other psychotic disorders. Schizophrenia disrupts multiple levels of language processing, from lower-level sensory processing to higher-level processing involved in inferring intent and understanding meaning(5,6). Neuroimaging studies have shown deficits in language pathways needed to process low-level perception and higher-level linguistic representations. A systematic review by Cavelti et al. identified a loss of gray matter in regions associated with language processing in adults diagnosed with schizophrenia (9). Deficits were found to be most prominent in the inferior frontal gyrus (involved in language processing and comprehension), the superior temporal gyrus (involved in auditory processing), and the inferior parietal lobe (involved in language comprehension and production) (10–14). These findings indicate that structural neurological changes underlie the inability for individuals with schizophrenia to effectively process and integrate language at all levels of linguistic representation.

Further neuroimaging studies are needed to correlate clinical characteristics of positive and negative thought disorder with specific deficits in these language pathways.

Binomial expressions are a frequently used linguistic feature in the English language, defined as pairs of words joined by a conjunction, usually “and” or “or.” Some binomials appear in a fixed order (e.g. ‘salt and pepper’ instead of ‘pepper and salt). These conventional orderings are shaped by abstract linguistic knowledge and direct experience (15–17). Abstract knowledge guides the processing of novel binomials, while direct experience influences the processing of frequently encountered binomials. Our previous work has shown that thought disorder is associated with the use of atypical binomial orderings in patients with psychotic disorders which is consistent with proposed links between thought disorder and disruptions in language processing at the syntactic and lexico-semantic levels (18,19). Therefore, binomials may serve as a quantifiable correlate of the underlying linguistic impairments characteristic of thought disorder.

We first sought to replicate two important findings from the literature. First, we aimed to replicate previous findings that given a pair of words, semantic and syllabic qualities of the word can accurately predict the preferred ordering in a large corpus of written works and in spontaneous speech (15,20). Second, we wanted to replicate our previous finding the binomial orderings in spontaneous speech are related to clinical scores for thought disorder in patients with psychotic disorders. Finally, we sought to expand on these findings by testing the hypothesis that increased use of atypical binomial orderings can distinguish cases and controls.

## Methods

### Participants and procedure

Patients with early course psychotic disorders (within 4 years of initial diagnosis of schizophrenia spectrum disorder, bipolar disorder with psychotic features, or psychosis NOS) were recruited from inpatient units and outpatient clinics at McLean Hospital. Control participants were recruited through online advertisements. Exclusion criteria were a history of major medical illness, severe head injury, hearing loss, or electroconvulsive therapy. Additionally, control participants had no history of psychiatric treatment or diagnosis. For demographic information about participants, see Table 1. Hospitalized patients had their capacity to consent independently evaluated by a psychiatrist unaffiliated with the project. All participants gave written informed consent for all study procedures. All study procedures were approved by the Institutional Review Board of Mass General Brigham. Patient diagnoses were confirmed with the Structured Clinical Interview for DSM-5 (21). All participants completed a video recorded interview. This interview included the Structured Clinical Interview for the Positive and Negative Symptom Scale as well as additional questions drawn from other clinical scales(22,23). PANSS scores were converted to factor scores using a well-established five factor model(24).

**Table 1.** Demographic and clinical characteristics of control and case samples. There were no statistically significant differences in age or sex between the groups.

|  | Controls | Cases |
| --- | --- | --- |
| **Sample Size** | 22 | 22 |
| **Age in years, mean (SEM)** | 24.6 (0.8) | 22.9 (0.7) |
| **Sex, N<sub>male</sub>/N<sub>female</sub>** | 9/13 | 8/14 |
| **Chlorpromazine Equivalents in mg, mean (SEM)** | 0 | 209.1 (47.9) |
| **PANSS** |  |  |
| **Positive Factor (SEM)** |  | 6.9 (0.5) |
| **Negative Factor (SEM)** |  | 9.8 (1.2) |
| **Excited Factor (SEM)** |  | 3.8 (0.2) |
| **Disorganized Factor (SEM)** |  | 4.5 (0.3) |
| **Depressed Factor (SEM)** |  | 6.5 (0.6) |

### Binomial analysis

The videorecorded interviews were transcribed by an expert transcriptionist who was not affiliated with the study and was blinded to the purpose of the study. In keeping with our previous work, we identified pairs of words linked by the word “and” (18). We rejected pairs that clearly crossed sentence boundaries and pairs where the “and” served to link two distinct phrases (for example, “I went to the store and Mark went to the zoo.”). To align with previous work, we also rejected pairs that used extender phrases (for example, “and stuff”) (20). The remaining phrases were considered binomials for all further analyses. For each binomial, we constructed a binomial with reversed ordering. For example, for the observed binomial “hat and shirt” we constructed the binomial “shirt and hat”.

We used the ngramr package in R to query the Google N-gram database for works published in United States English between 2010 and 2019 (25–27). For each binomial, we calculated the ratio of the occurrences of alphabetical ordering in this database to the occurrences of the either the alphabetical or the reverse alphabetical ordering. We called this ratio the attested ordering.

We used a logistic regression model created by Morgan and Levy (2006) to calculate an ordering preference based on abstract features(15). These authors identified seven features that predicted binomial ordering in both a large corpus and in experiments with healthy control participants(15,16). The seven features are formal markedness, perceptual markedness, power, iconic or scalar sequencing, final stress, frequency, and length(15). For full details of these features, see (15,17,20). Formal markedness means that the word with the broader or more encompassing meaning comes first. For example, in “pull and yank”, pull is more general than yank. Perceptual markedness includes multiple constraints such as animate before inanimate and concrete before abstract. For example, in “goats and bushes”, goat is animate and so comes first. Power means that the more culturally powerful word comes first. For example, in “kings and beggars”, kings are more powerful and so come first. Iconic or scalar sequencing means that things that happen in sequence are ordered in that sequence. For example, in “start and finish”, start precedes finish temporally and so that ordering is preferred. Final stress means that the final syllable of the second word should be unstressed if possible. For example, in “guitar and photography”, the final syllable of guitar is stressed while the stress is on the second syllable of photography and therefore guitar should come first. Frequency means that the more frequent word should come first. We calculated frequency from the same Google N-gram database. For example, in “hiking and “skiing”, hiking appears more frequently than skiing and so should come first. Length means that the word with fewer syllables should come first. For example, in “guitar and photography”, guitar has two syllables and photography as four and therefore guitar should come first. For each feature and each binomial, two authors (AL and MM) independently scored whether that feature favored the alphabetical ordering, the reverse alphabetical ordering, or was not active. For each feature and each binomial, we generated a consensus score. We then used the regression coefficients from the Morgan and Levy paper to generate an estimate of the likelihood of the alphabetical order based on our scores. We called this likelihood the abstract ordering. Finally, again using regression coefficients from Morgan and Levy we combined the attested and abstract ordering to generate an overall ordering likelihood(15).

### Statistics

Unpaired t-tests were used to compare controls and patients. Spearman correlation was used to measure the relationships between clinical features and linguistic measures. Multiple comparisons were addressed using the Dubey-Armitage-Parmar correction (28).

## Results

### Abstract features and attested ordering make similar predictions about spoken binomial ordering preferences

Across all interviews, we identified 322 unique binomials, 43 of which were not found in the Google n-gram corpus and therefore did not have an attested order. In previous reports, abstract order and attested order were highly, but not completely, correlated (15). For the 279 binomial pairs with attested orderings, we found that the abstract ordering calculated from the Morgan and Levy regression coefficients was highly correlated with the attested ordering observed in the Google n-gram corpus (Figure 1a, Pearson’s R = 0.72, p<10^−4^). Out of the 279 binomials with both attested and abstract ordering values, only 39 had opposing preferences between attested and abstract. Out of these binomials, the average difference in preference strength was 0.35 which indicates that this occurred mostly for binomials that had moderate ordering preferences rather than strong ordering preferences. Overall predicted ordering was calculated using a weighted average of the abstract and attested ordering (or only the abstract ordering for those binomials that did not appear in the Google corpus). For all binomials, the strength of the predicted ordering was clearly non-normal (Chi-squared goodness of fit test p<10^−5^) and non-uniform (Chi-squared goodness of fit test p<10^−19^, Figure 1b). Compared to a random distribution of ordering preference strength, more binomials had a strong predicted ordering preference (z-score of .975-1.0 bin was 3.96., corresponding to Bonferroni adjusted p<0.002).

**Figure 1.**
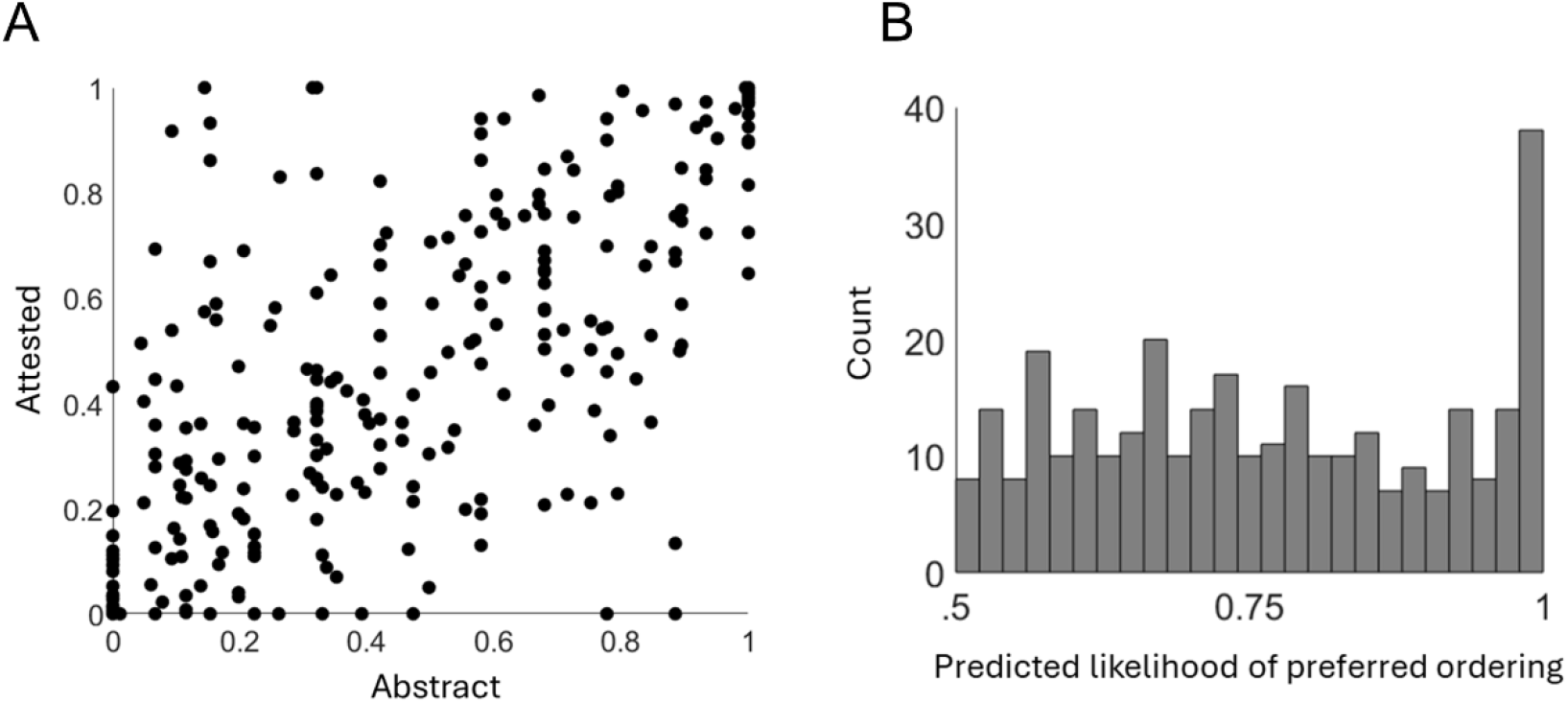
Abstract and attested binomial ordering predictors. **A**. Scatter plot of the calculated abstract ordering preferences for each binomial vs their ordering in the Google n-gram corpus (“attested”). **B**. Histogram of the strength of the preferred ordering where 0.5 indicates no preference between the alphabetical and reverse alphabetical ordering.

### Atypical binomial orderings correlate with case-control status and thought disorder symptoms

For each participant, we calculated the mean likelihood of their binomial orderings. Patients had statistically significantly lower mean likelihoods (0.61 SEM 0.03) compared to controls (0.72 SEM 0.02, p = .0025, unpaired t-test, Cohen’s d = 1.0, Figure 2a). There was no relationship between case/control status and the likelihood of using non-attested binomials (unpaired t-test, p = .24). In line with our previous work, we consider any ordering with a predicted likelihood of less than 0.33 to be a “rare” ordering (18). For each participant, we calculated what proportion of their binomials were rare orderings (Figure 2b). There were no statistically significant relationships between the total number of binomials uttered and the proportion of rare orderings in either patients (p >.3) or controls (p>.12). Patients had statistically significantly higher proportion of rare orderings (0.18 SEM 0.04) compared to controls (0.06 SEM 0.02). We previously showed that the proportion of rare orderings was associated with thought disorder. Here, we replicated this finding and found that, in patients, the Disorganization factor was statistically significantly associated with the proportion of rare orderings (Figure 2c, Spearman’s rho = 0.53, p_adj_ = 0.039). There were no statistically significant relationships between rare ordering and the other symptom factors (all p_adj_ > 0.7) nor was there any relationship between rare ordering and chlorpromazine equivalents (p > 0.09).

**Figure 2.**
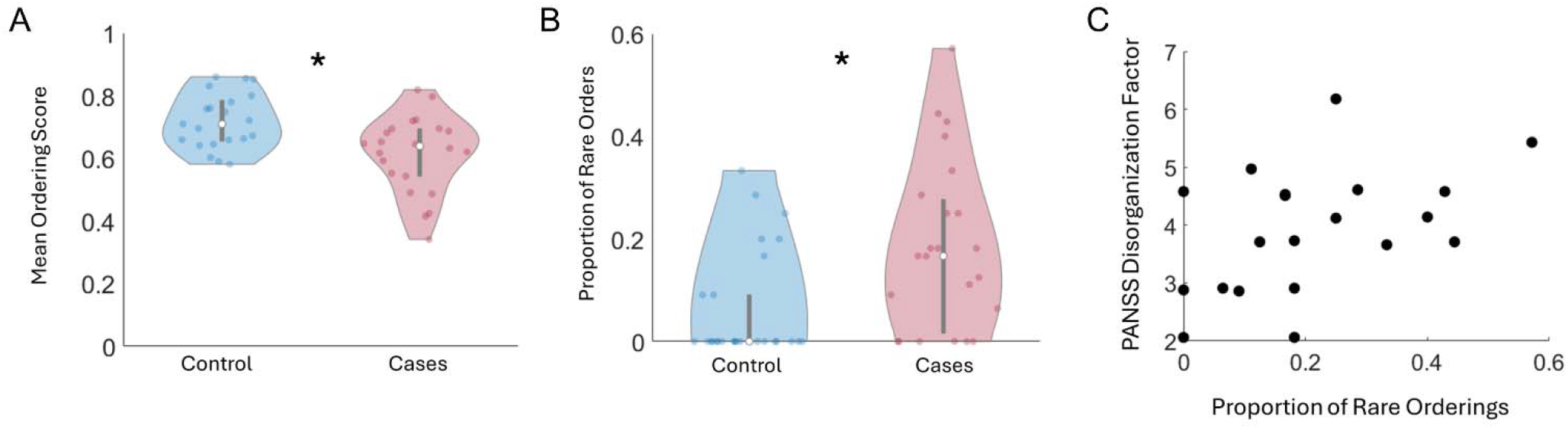
Binomial orderings in cases and controls. **A**. Violin plots of the mean likelihood for binomial orderings uttered by controls (blue) and cases (red). * p <.003 unpaired t-test. **B**. Violin plots of the proportion of uttered binomials that were in a rare ordering for controls (blue) and cases (red). * p <.003 unpaired t-test. **C**. Scatter plot of the relationship between the proportion of uttered binomials that were in a rare ordering and the PANSS Disorganization factor in patients only.

## Discussion

We analyzed the ordering of spontaneously uttered binomial phrases in patients with psychotic disorders as well as healthy controls. In keeping with previous work, we found that a prediction model using semantic and linguistic features of the binomials to predict ordering was correlated with ordering preferences observed in a large corpus. We found that patients were more likely to violate the predicted orderings compared to healthy controls. While some binomials have only weak ordering preferences, we found that the differences between cases and controls were likely related to the increased use of highly disfavored orderings in patients. Furthermore, we found that use of these rare orderings was related to thought disorder symptoms and not other symptoms domains or psychiatric medications.

### Thought disorder and atypical binomial orderings

Despite showing a preference for attested binomials with strong conventional orderings, individuals with psychosis demonstrated more disorganized language patterns in our study, as evidenced by their increased use of atypical binomial orderings. Given that there was no relationship between the number of binomials used and the percentage of rare binomial orderings, it is likely that the increased use of atypical binomials in individuals with psychosis is not due to increased language output but rather due to dysregulated language processes. Research by Morgan and Levy found that direct experience plays a significant role in the processing of frequently encountered binomials (15). Instead of being processed as individual components, these binomials are stored in memory as complete phrases and are recalled holistically, making processing more efficient. The more often a binomial is encountered, the faster it is processed, with a stronger and more consistent preference for the familiar word order (15,29). This may explain the preference for binomials with strong ordering preferences seen in both individuals with psychosis and controls. However, in individuals with psychosis, cognitive processes may be disrupted, leading to inconsistent or disordered recall of these patterns and the inability to reliably produce these conventional orderings

The use of atypical binomial orderings in individuals with psychosis can be linked to cognitive and neurological deficits. Marini et al. found that while individuals with schizophrenia may experience impairments in sentence-level processing, their primary challenge lies in maintaining narrative coherence, which is dependent on executive function (30). Executive function depends on top-down processing, which enables individuals to use higher-order cognitive control to integrate meaning and structure by coordinating goal-directed behaviors and regulating attention and inhibition. Executive function may help produce conventional orderings for rarely encountered binomials. The role of top-down processing is supported by Fedorenko et al., who found that the fronto-temporal language network is globally activated for both syntactic and semantic processing, without any region specifically dedicated to either function (31). Efficient language production relies on integrated processing across these networks. When this integration breaks down, individuals with psychosis may shift towards bottom-up processing, as they struggle to use top-down predictions based on prior experience (5). This could lead to increased randomness and therefore higher frequency of atypical binomial orderings seen in our results.

Thought disorder may be tied to the degree of disruption in the fronto-temporal network, with more severe breakdowns leading to increased disorganization symptoms. Given that disorganized language patterns may reflect underlying cognitive and neurological deficits, tracking these changes in language output could offer an accessible way to monitor and predict the onset and progression of psychosis. Studies on symptom trajectories in psychosis have shown that persistent disorganized language use is a significant predictor of psychosis onset (8,32). Early identification of formal thought disorder through atypical binomial use can therefore be a potential avenue for early intervention for individuals at high risk for developing psychotic disorders

### Limitations and future directions

While this study is an advancement on previous work, it has several limitations that should be considered when interpreting our findings. This is a cross-sectional study and future longitudinal studies should establish whether atypical binomial orderings are a trait marker of psychotic disorders or if they are sensitive to current symptom burden. Thought disorder in this study was assessed as a unitary entity using the PANSS. Thought disorder in clinical practice encompasses several distinct subtypes which are not captured in the PANSS Disorganization factor. For example, negative thought disorder (decreased speech and paucity of thought) is not included in this factor. However, some of these elements are included in other the symptom factors which were not found to be associated with binomial ordering.

Therefore, our results suggest that binomial ordering is a marker of positive thought disorder, but rigorous studies using thought disorder specific scales are needed to fully establish this.

## Acknowledgements

This work was supported by K23MH118565 from the National Institute of Mental Health (MM).

## Author Contributions

AL wrote the manuscript and analyzed data, JL collected data and edited the manuscript, MM wrote the manuscript and analyzed data.

## Data Availability

Binomial lists for all participants are available upon request.

